# Flowable Grafts Made from Granular Extracellular Matrix (gECM) Hydrogels Promote Integrative Repair of Articular Cartilage in a Large-Animal Model

**DOI:** 10.64898/2026.05.05.723111

**Authors:** Jeanne E. Barthold, Juliet Heye, Kaitlin P. McCreery, Lea Savard, Katie Bisazza, Emily Miller, Hongtian Zhu, Woowon Lee, Maxwell C. McCabe, David Ceja Galindo, Shannon Blanco, Virginia Ferguson, Nancy Emery, Brian Johnstone, Benjamin Gadomski, Stephanie E. Schneider, Jeremiah Easley, Corey P. Neu

**Author notes:** Corresponding author: Corey P. Neu, 1111 Engineering Drive, UCB 427, Boulder, CO, 80309, USA.

## Abstract

Focal injuries to articular cartilage in load-bearing joints fail to heal and often progress to degeneration, underscoring the need for repair strategies that result in restored cartilage structure and function rather than fibrocartilage formation. Granular extracellular matrix (gECM) hydrogels, flowable grafts composed of densely-packed matrix particles, offer a promising approach but lack long-term functional validation in large-animal models. Here, we developed a flowable gECM hydrogel composed of decellularized cartilage microparticles incorporated within a thiol-functionalized hyaluronan matrix. Proteomic analysis confirmed enrichment of cartilage-specific gECM matrisome components. When implanted into critical-sized femoral condyle defects in a goat model and evaluated 12 months post-implantation, both gECM hydrogel and microdrilling (surgical controls) achieved >80% defect filling. However, in contrast to microdrilling, gECM repair tissue exhibited surface tribological (friction, adhesion) and compressive mechanical properties comparable to native cartilage, with a similar proteoglycan-to-collagen ratio, enrichment of type II collagen, minimal type I collagen (typical of a fibrous scar), improved quantitative MRI metrics, and evidence of lateral cartilage integration and subchondral bone remodeling. Together, these findings demonstrate that a flowable gECM hydrogel supports integrative, cartilage-like repair in a load-bearing joint, supporting advancement of this approach toward clinical translation.

**One Sentence Summary:** A granular ECM hydrogel implanted in a goat condyle provided a robust repair, filling the defect tissue with integrated, hyaline-like cartilage at 12 months.

## INTRODUCTION

Knee pain is a serious and widespread societal problem that affects approximately 25% of adults, limiting mobility and impacting quality of life (*1*). The most common cause of knee pain is osteoarthritis (OA), a degenerative joint disease that is difficult to treat and associated with chronic pain and disability. In 2019, an estimated 528 million people worldwide were living with OA (*2*). The degeneration processes that lead to OA can be triggered by acute events (e.g. mechanical insult, focal cartilage damage, or adjacent injury such as anterior cruciate ligament rupture) and are characterized by a cascade of inflammatory events in the joint that result in progressive cartilage degradation.

Natural, non-interventional repair of cartilage defects is uniquely challenging because articular cartilage is an avascular tissue with limited intrinsic regenerative capacity (*3*). Microdrilling is a commonly used surgical treatment for focal cartilage defects and remains practiced as a single-stage procedure, despite limited evidence supporting long-term efficacy (*4*). The procedure involves creating small perforations in the underlying subchondral bone beneath a cartilage lesion to stimulate a repair response that recruits marrow-derived progenitor cells, resulting in defect filling with a blood clot. Clinical problems with this historical standard-of-care procedure are well known, including repair tissue that is predominantly fibrocartilaginous and that fails to mimic the structure, function, and durability of native hyaline cartilage. Microdrilling can also increase surface roughness and friction within the joint. Importantly, microdrilling has not been shown to meaningfully delay or reverse long-term joint degeneration. The most effective current clinical treatment for cartilage defects involves viable osteochondral allografts; however, this approach is limited by donor tissue availability and the short storage time of living grafts (approximately 30 days).

Successful cartilage repair and regeneration remain significant unmet needs in medicine. Granular hydrogels represent a relatively recent innovation in tissue engineering that may provide an ideal material composition and advantageous platform for cartilage repair. Cartilage presents distinct biological and mechanical demands, as it must support cell viability and matrix maintenance in an avascular environment while simultaneously withstanding high compressive and shear forces during everyday activities like standing, walking, and jumping. Granular hydrogels offer a modular design composed of densely packed particle inclusions, often combined with a biopolymer base. The source, size, and packing density of the inclusions, as well as the properties of the hydrogel component, can be tailored to meet tissue-specific demands. Recent work has shown that rod-like particles promote more cellular invasion and activity compared to spherical particles (*5*). Moreover, when inclusions are densely packed, the bulk mechanical behavior of the granular hydrogel is dictated by the mechanics of the stiffer particles, rather than those of the surrounding hydrogel (*6*).

Although a variety of granular hydrogels have been evaluated *in vitro* and in short-term or small-animal *in vivo* studies, none have been evaluated in long-term, functional studies relevant to cartilage repair (*5, 7–15*). Understanding long-term tissue integration, durability, and mechanical function is essential for translational success, as many previous candidate cartilage therapies have demonstrated promising early outcomes but failed to produce sustained improvements in healing or pain beyond six months (*16*). Additionally, evaluation of potential treatments in large-animal models at clinically-relevant defect sizes is necessary for translation to human clinical trials and commercial therapies.

To address the need for cartilage repair materials that support biological function while also withstanding physiological mechanical loading, we developed a novel and flowable granular extracellular matrix (gECM) hydrogel composed of cartilage-derived matrix particles sourced by grinding and decellularizing cartilage tissue. The particles are densely packed within a thiol-functionalized hyaluronan hydrogel that undergoes spontaneous, thermally-dependent crosslinking upon mixing, delivery, and exposure to body temperature. The gECM hydrogel exhibits mechanical robustness due to dense particle packing. We have previously explored simple proof-of-concept bioprinting using cartilage particles combined with hyaluronan (*17*). However, this material was generated using harsh processing, chemical agents, and uncontrolled formulations unsuitable for translation. Moreover, defined material compositions with particle sizes suitable for translation were largely undefined, and *in vivo* efficacy was unexplored. Here, motivated by benchtop characterization and refined biofabrication approaches, we evaluate the long-term efficacy of a gECM hydrogel in a critical large-animal study designed to assess the translational potential for cartilage repair.

## RESULTS

### Granular ECM particles were decellularized, but maintained structural composition similar to native cartilage tissue

Granules were sourced from porcine hyaline cartilage tissue and decellularized using an SDS rinsing protocol (**Figure 1A**). The particles were rendered decellularized, as they contained <50ng dsDNA/mg of tissue (*18*) (**Figure 1B**). Native tissue and decellularized granules were characterized for protein composition using mass spectrometry specialized for extracting insoluble extracellular matrix proteins as well as soluble factors (**Figure 1C-D, Supplemental Figure 1**). The processing of the granules resulted in a three-fold reduction of nuclear proteins (3.2% in native tissue to 1% in granules), membrane proteins (2.2% to 0.6%), and a 9-fold reduction in cytoskeletal proteins (4.4% to 0.5%), all percentages of total label-free quantification (LFQ) signal, supporting our observation of effective decellularization. Matrisome proteins represented 89.8% of total detected signal for native tissue, compared to 97.9% of total signal intensity in decellularized granules. Matrisome proteins have been classified into two primary categories, each of which is further divided into three functional subcategories: core matrisome proteins (including collagens, proteoglycans, ECM glycoproteins) and matrisome-associated proteins (including ECM-affiliated proteins, secreted factors, and ECM regulators). The large majority of the matrisome was composed of core matrisome proteins (97% of matrisome protein signal in both native tissue and decellularized granules). The largest matrisome subcategory difference observed in the granules after processing was a change in proteoglycan percentage from 17.5% in native matrisome to 6.42% in decellularized matrisome. This finding implies that the processing removed proteoglycans, though this was expected based on previously reported decellularization methods (*19, 20*). However, collagen concentration maintained the majority share of the granules at 86% of total LFQ signal, while ECM glycoprotein concentration was reduced by only 50% through processing (from 7.5% to 3.7%). COL2A1 was the most abundant protein identified in both native and decellularized tissue. Through processing of the tissue into granules, all matrisome-associated subcategories were reduced by at least 6-fold and make up less than 0.5% of detected signal within the granule matrisome (secreted factors were 0.04%, ECM-affiliated proteins were 0.13%, and ECM regulators were 0.16% of total matrisome protein signal).

**Figure 1.**
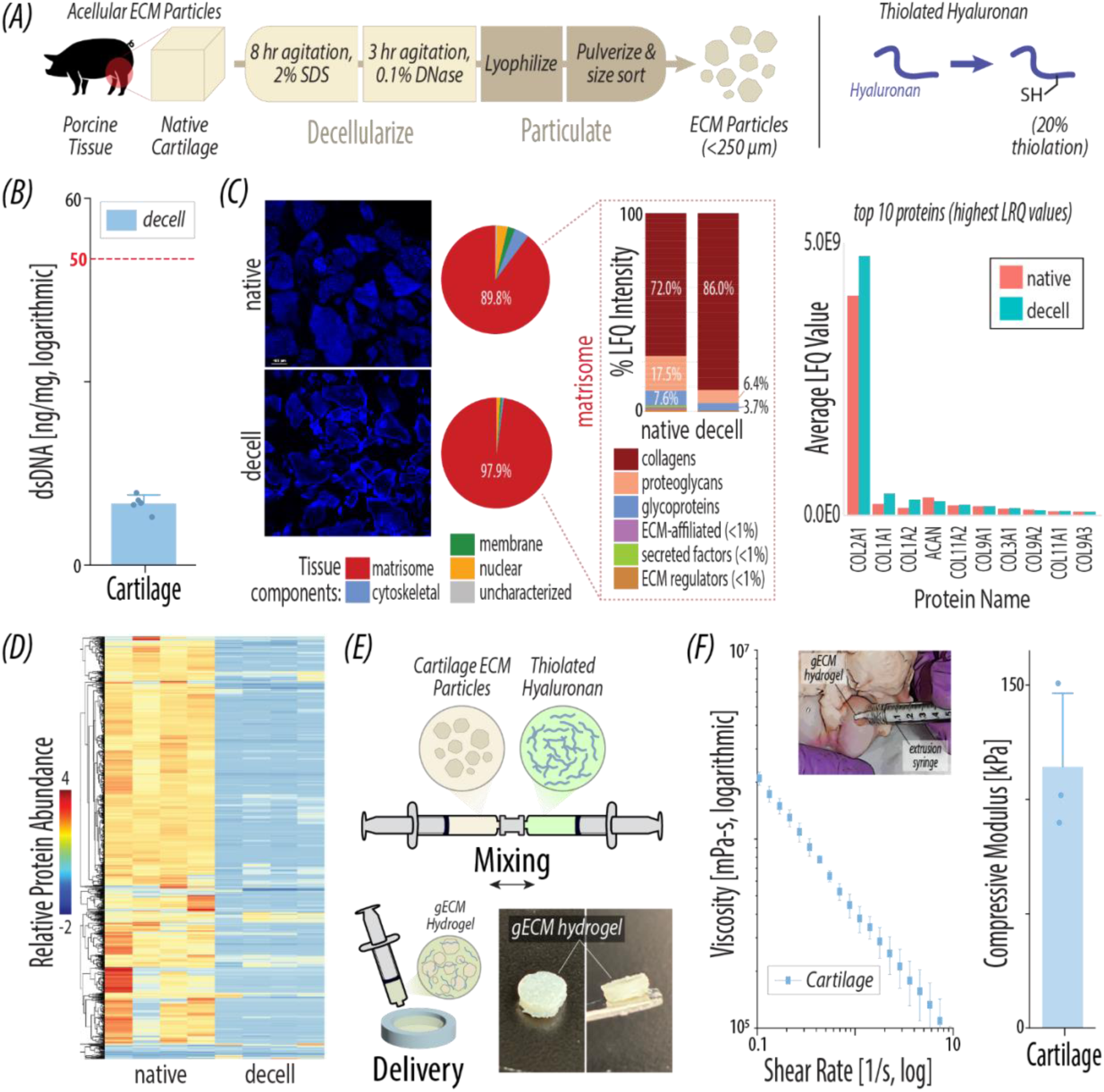
Granular extracellular matrix (gECM) hydrogels were manufactured from particulated animal tissues in combination with functionalized macromolecules. *(A)* ECM granules sourced from porcine articular cartilage maintain key structural proteins after decellularization and, despite removal of some proteoglycans, secreted factors, and ECM regulators, produce a robust mechanical scaffold when combined with thiol-modified hyaluronan. Tissue was extracted from porcine knee joints, decellularized, rinsed, and pulverized into lyophilized granules prior to combination with a thiol-modified hyaluronan solution. *(B)* Particle decellularization is confirmed using a genomic DNA miniprep kit. *(C)* Mass spectrometry performed on porcine granules pre- and post-decellularization revealed matrisome proteins, specifically collagens, as the most highly conserved components. The most abundant proteins in the final processed granules by LFQ intensity are all collagens, dominated by COL2A1. *(D)* Heat map of all identified proteins (left) demonstrating relative abundance throughout decellularization (left four columns native, right four decellularized). *(E)* To form stable 3D constructs, decellularized granules in saline were mixed with functionalized (20% thiol-modified) hyaluronan. Following mixing, gECM hydrogels were delivered in various shapes through flowing and extrusion. *(F)* gECM hydrogels demonstrated shear-thinning behavior and flowability immediately post-mixing. Following 30 minutes of polymerization at 30 °C and 12 hours in standard saline, bulk compressive modulus of benchtop and acellular gECM hydrogels slightly exceeded 100 kPa, representing the material stiffness expected at the time of initial implantation.

### gECM granules combined with thiol-modified hyaluronan form stable gECM hydrogels

We have previously shown that 250 µm ECM granules (gECM) packed tightly into a PEGDA/tHA (thiol-modified hyaluronan) gel resin form a stable 3D scaffold with mechanical properties under compressive loading approaching the values of native cartilage (*6*). Additionally, we previously showed that 100 µm gECM can be combined with thiolated hyaluronan alone and can be extruded into stable 3D constructs via direct ink writing 3D printing (*17*). Here, we validated that when 250 µm granules are mixed at a high density (0.2 g/mL) with thiolated hyaluronan alone (without PEGDA or other external crosslinkers), the material is flowable during delivery, exhibiting shear-thinning behavior, but then also crosslinks into a 3D structure at 37 °C (**Figure 1E-F**). Furthermore, the crosslinked construct demonstrates compressive modulus values similar to those made with tHA/PEGDA crosslinking. Under compression, at particle density values just below the previously identified percolation threshold (0.57 v/v) (*6*), the gECM hydrogels maintain a modulus of ∼114 kPa.

### Goats returned to normal function immediately following gECM hydrogel implantation and recovery from anesthesia

Inspired by our finding that gECM hydrogels are flowable but crosslink at body temperature following extrusion and delivery, and further by the concept of rebuilding a tissue as a sum of its particles, we evaluated the performance of gECM hydrogels in a large-animal (caprine) model of cartilage defect repair (**Figure 2A**). Goats were placed under general anesthesia and, in bilateral condyles, critical-sized defects (6.5 mm diameter × 2.5 mm depth) were placed in the medial femoral condyles of contralateral joints. The base of the defects was drilled with 1 mm drill bits to create five holes, following a standard microdrilling approach, which has shown superior patient-reported outcomes and lower revision rates compared to microfracture (*21*). The defects were then either left empty (control) or augmented with delivery of gECM hydrogels (**Figure 2B**). Within 5 minutes of mixing, the gECM hydrogel was delivered by flowing into the defect and sculpted by the surgeon to match the contours and shape of the cartilage surface. After surgery and recovery from anesthesia, goats were fully weight-bearing within 45 minutes and underwent normal activities over 12 months to allow for evaluation of a natural healing response to implants (**Figure 2C**). Goats were evaluated throughout the 12 months with periodic radiographs which showed no observable joint space narrowing or obvious bone pathology (**Supplemental Figure 2**).

**Figure 2.**
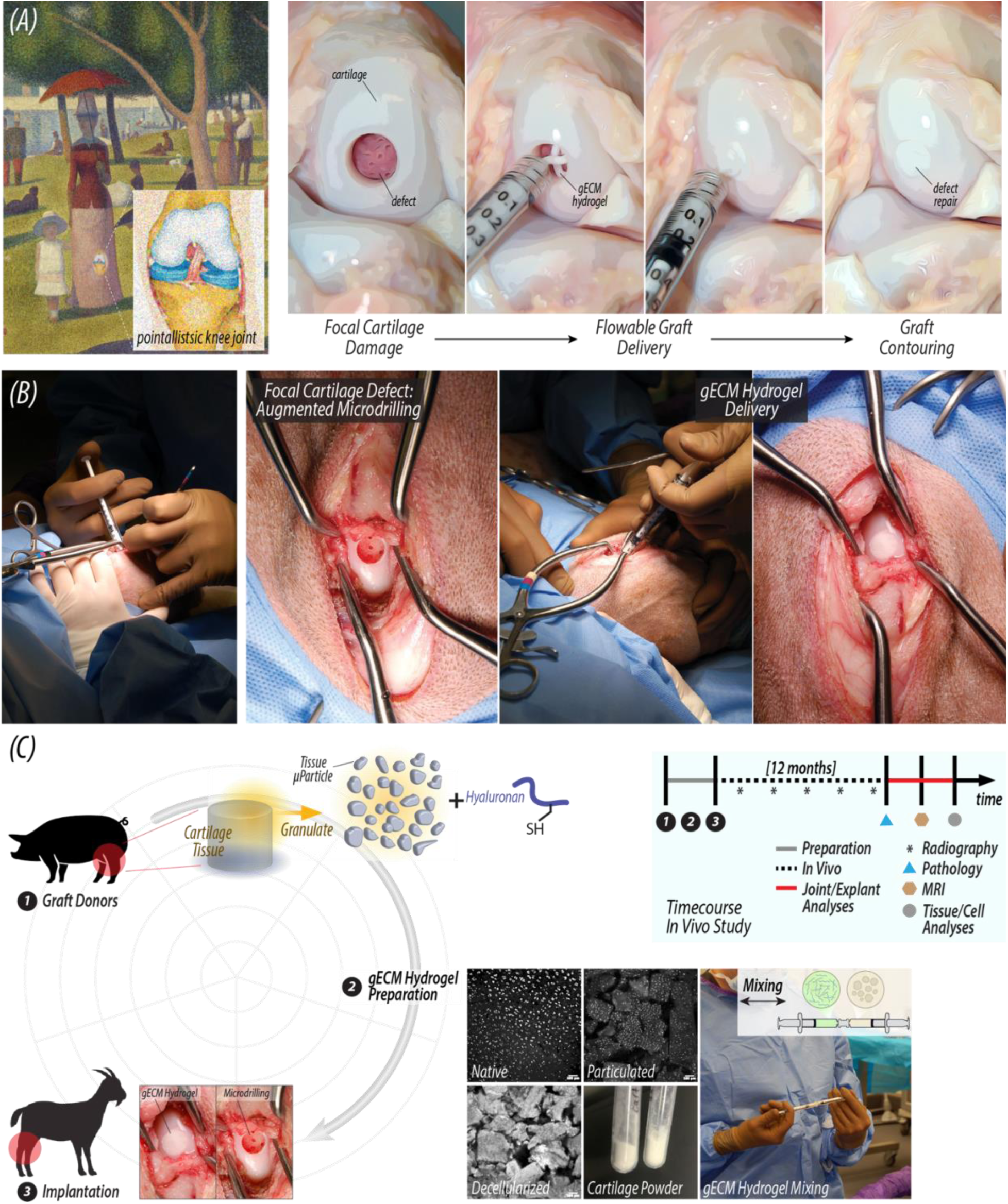
gECM hydrogels were evaluated in a 12-month, large animal cartilage defect repair model, against microdrilling as a standard of care. *(A)* Inspired by our finding that gECM hydrogels are flowable but crosslink at body temperature following extrusion and delivery, and further by the concept of rebuilding a tissue as a sum of its particles (analogous to pointillistic paint patterns in Georges Seurat’s *A Sunday Afternoon on the Island of La Grande Jatte*), we transitioned to evaluate performance of gECM hydrogels in a large-animal (caprine) model of cartilage defect repair. Here, we use cartilage granules to reconstruct a tightly-packed 3D construct that restores the cartilage as a single whole. Flowability of the gECM hydrogel enables custom and real-time filling of a focal cartilage defect that can be precisely contoured prior to polymerization at body temperature. *(B-C)* We utilized our flowable gECM hydrogel to fill a critical-sized goat (caprine) condyle. Contralateral limbs were left unfilled, treated with microdrilling alone but additional augmentation or material fill. Goats were immediately weight bearing following recovery from anesthesia, and unrestricted in movement for 12 months. Multiple assessments of joint and animal health were performed prior to and following euthanasia, including radiography pathology, MRI, and tissue/cell level analyses.

### Twelve months following implantation, cartilage defects remained filled with hypercellular repair tissue without evidence of gross systemic pathology

After euthanasia at 12 months, visual inspection following dissection showed that defects of most joints – both microdrilling (control) and gECM (treated) joints – demonstrated >80% tissue filling (**Figure 3, Supplemental Figure 3**). Tissue fill was less variable for gECM hydrogels (**Figure 3C**), and interfacial regions between the repair and surrounding tissue (dashed lines in **Figures 3A-B**) showed evidence of continuity between tissues without obvious gaps or residual defects that may promote mechanical instabilities (e.g., stress concentrations at regions of constricted geometry). Microdrilling and gECM hydrogels showed increased cell density compared to native cartilage (p<0.0001). Additionally, histological data was used for standard metrics in cartilage repair, including ISO 10993 scoring, Modified O’Driscoll’s Score, and the International Cartilage Repair Society (ICRS) II score (**Supplemental Figure 4**). Furthermore, we did not measure significant differences between treatment groups in any of the evaluated criteria (p>0.05). We also did not find evidence of substantial gross pathology, with normal appearance of lymph nodes, lung, liver, heart, kidney, spleen, and synovium (**Supplemental Figure 5**). Because no differences were found in surgical outcomes, observable tissue filling, histological scoring, and gross pathology, we turned next to evaluate the mechanical function and detailed composition of infilled tissues.

**Figure 3.**
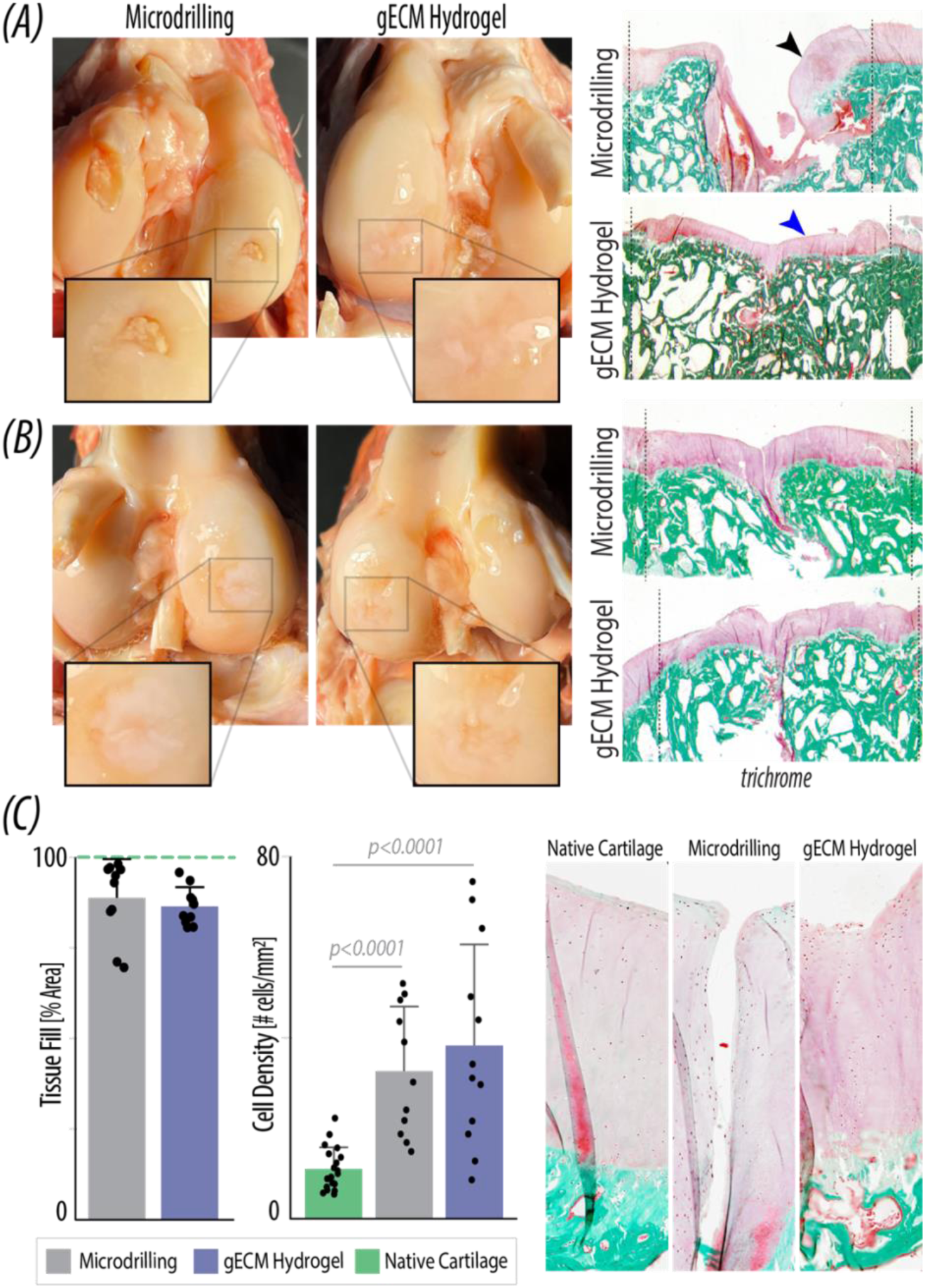
12 months following implantation, cartilage defects remained filled with hypercellular repair tissue. *(A-B)* Representative macro images (photographs) and histological sections (trichrome staining) of both microdrilling and gECM hydrogel (as an augmented microdrilling procedure) contralateral limbs. Pink denotes cartilage while blue/green denotes bone. Dashed lines represent the margins of the defect during surgery where tissue has since infilled, with the interface between repair and surrounding cartilage showing largely continuous structural integration. Some joints demonstrated a lack of filled volume (black arrow), while other joints demonstrated structural filling over 12 months (blue arrow). *(C)* Microdrilling and gECM hydrogels demonstrated >80% tissue filling, though gECM hydrogels showed a less variable filling response. Microdrilling and gECM hydrogels also showed increased cell density (hypercellularity) compared to native cartilage (p<0.0001).

### gECM hydrogels restore low surface friction, adhesion, and roughness to values near native cartilage

The articular surface of cartilage exhibits tribological and compressive properties that support low friction and adhesion while also transmitting forces essential for healthy joint function (*22, 23*). Atomic force microscopy (AFM) (*24*) was performed on the tissue to evaluate surface properties of the cartilage and to compare responses of native tissue to the surface of gECM hydrogel or microdrilling repair tissue (**Figure 4A-B**). Analysis was performed after euthanasia and joint dissection to remove the repaired region with the surrounding native tissue. gECM hydrogel maintained lower levels of surface friction, adhesion, and roughness values compared to the surface of microdrilling cartilage. The friction coefficient at the surface of the gECM hydrogel repair was 0.22, similar to the native cartilage surface and comparable to historic values (*24*), and lower than the frictional coefficient of the microdrilling repair of 0.31 (p=0.071). Additionally, the gECM hydrogel surface measured adhesion of 3.58 nN, lower than the native cartilage surface, and lower also compared to microdrilling repair surface adhesion of 5.59 nN (p=0.081). Finally, the surface roughness of gECM hydrogel measured 346.42 nm, similar to the native cartilage surface, while the surface of the microdrilling repair had an elevated roughness of 431.72 nm.

**Figure 4.**
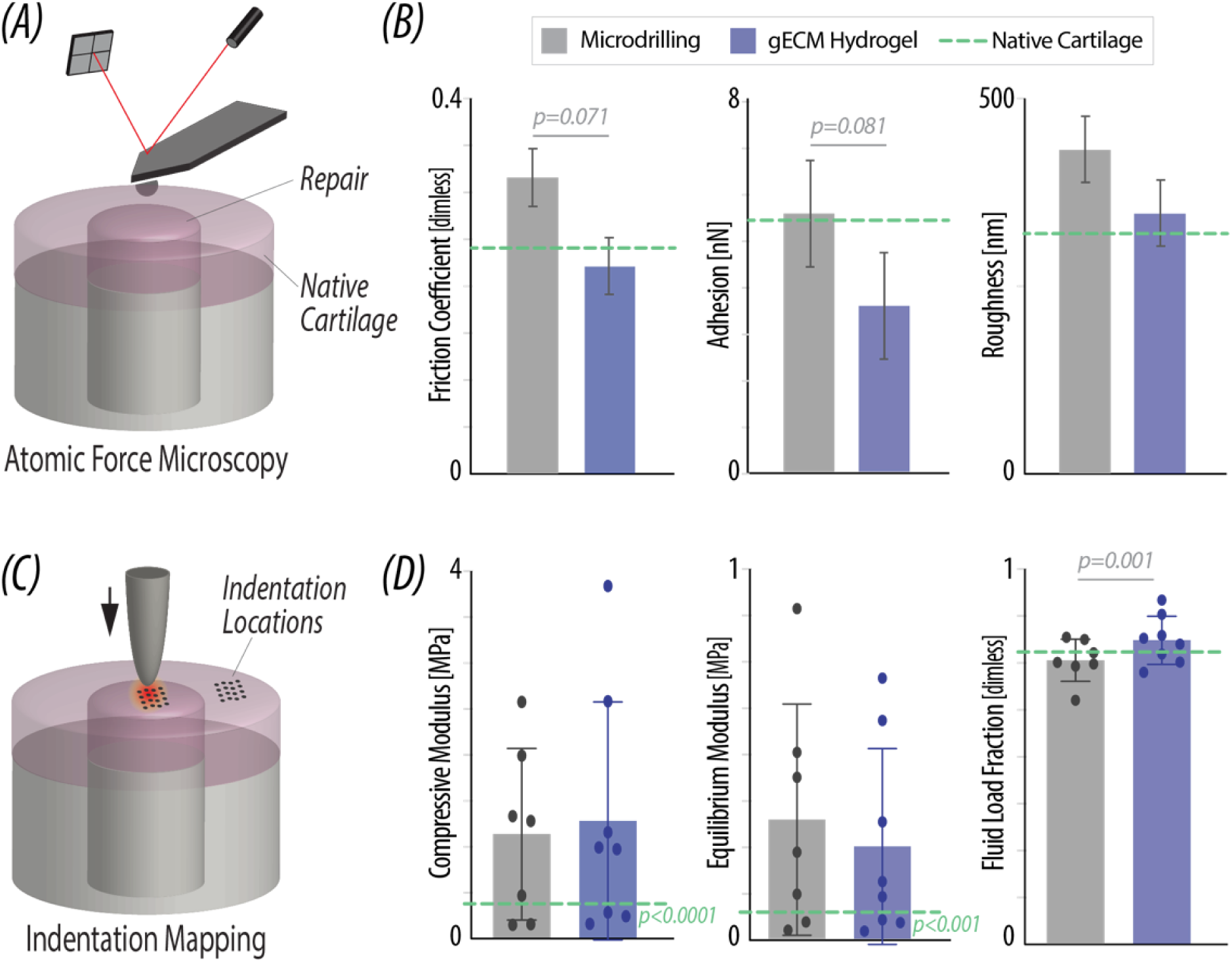
gECM hydrogels demonstrated low surface friction, adhesion, and roughness. *(A)* Atomic Force Microscopy was used on the surface of all repair regions to evaluate critical cartilage tribological measurements, *(B)* The gECM hydrogel (blue) maintained low frictional coefficient, adhesion levels, and surface roughness compared to microdrilling repair tissue (gray), and the surrounding native tissue (green dashed line). *(C)* Microindentation was used on the surface of all repair regions to further evaluate mechanics under compression, equilibrium, and the fluid load fraction of the tissue. *(D)* Compressive modulus and equilibrium modulus were significantly elevated in both treatment groups, compared to native tissue, while fluid load fraction did not significantly differ from native to treatment in either treatment. (p > 0.05, green p-value represents a significant difference of both repair types compared to native cartilage).

We further measured compressive properties at the articular surface using microindentation testing, enabling comparison of bulk biomechanical (indentation) properties (**Figure 4C-D**). We found that the compressive modulus of the gECM hydrogel was greater compared to microdrilling repair tissue (p=0.033), and both repairs were higher than the average native cartilage values (p<0.001). The equilibrium modulus of both repair types was also higher compared to native cartilage (p<0.001). The fluid load fraction of gECM hydrogel repairs was higher compared to microdrilling repair tissue (p=0.001), but both repairs were not different from native cartilage.

### gECM hydrogel showed a proteoglycan-to-collagen ratio similar to native cartilage, with enrichment of type II collagen, and minimal type I collagen that is typical of a fibrocartilaginous repair

We next turned our attention to examine the quality and composition of repair tissue filling the defect space, evaluated using immunohistochemistry and Raman spectroscopy. Articular cartilage is unique in its composition from other connective tissues, rich in predominantly type II rather than type I collagen. We performed immunostaining on tissue slices from repaired joints and found that the gECM hydrogel was rich in type II collagen and largely void of type I collagen (**Figure 5A-B**). Even at interface regions between the repair and host cartilage, we did not visualize type I collagen content, typical of fibrocartilaginous tissue, that often results as a common outcome in cartilage repair. In contrast, microdrilling repair tissue showed a mix of type II and type I collagen (red arrows), indicating the collagen content showed evidence of a more fibrous repair with elevated staining for type I collagen.

**Figure 5.**
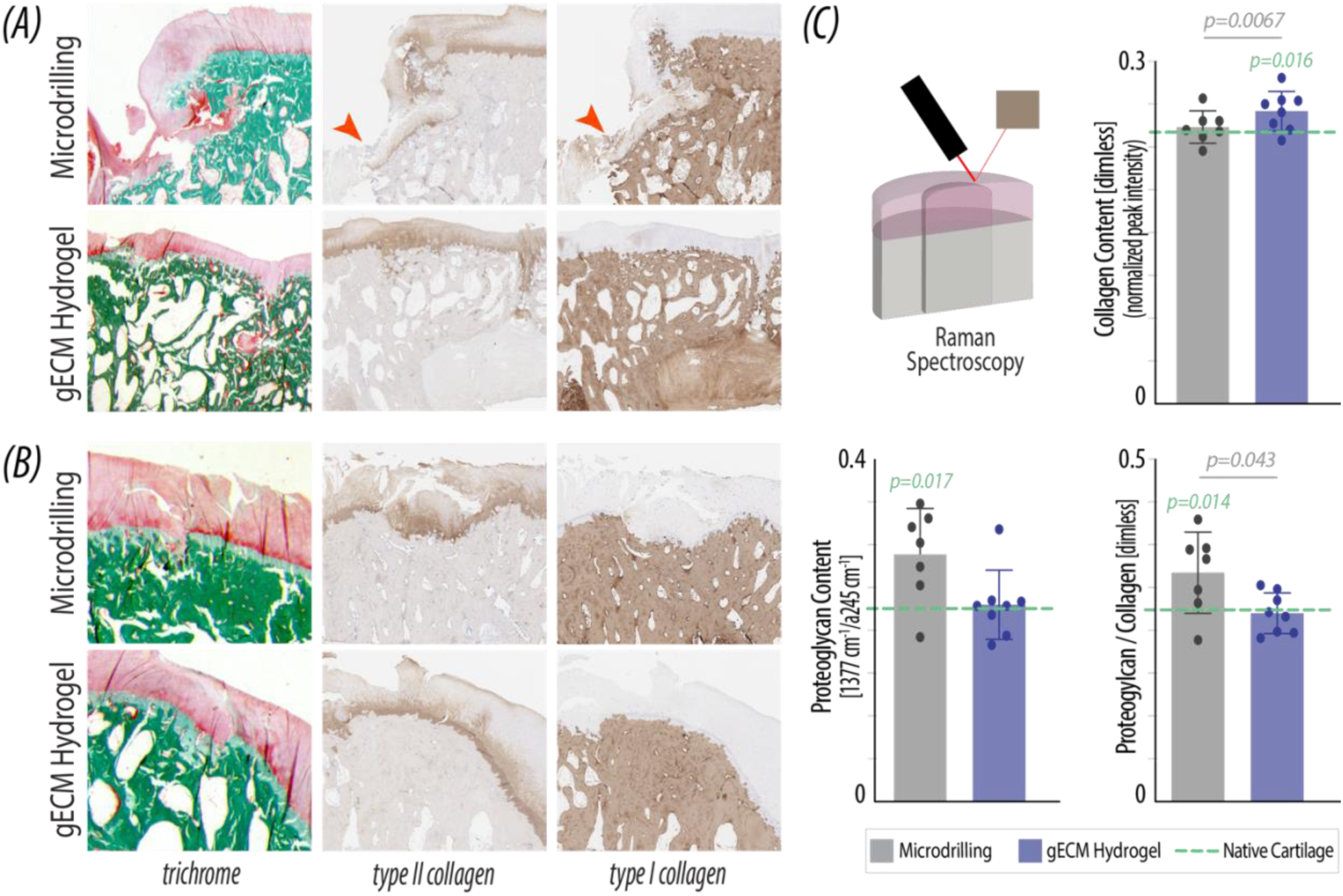
gECM hydrogel repair restores collagen II and proteoglycan content similar to that of hyaline cartilage. *(A-B)* Representative staining in two joints indicates presence of types II and I collagen, with type II collagen visualized throughout gECM hydrogels, and evidence of type I collagen in some microdrilling repair tissue (red arrows). *(C)* Raman spectroscopy quantified proteoglycan and collagen content of repair regions compared to the levels of surrounding native tissue (green dashed line). gECM hydrogel restored the native proteoglycan-to-collagen ratio, while in microdrilling repair this ratio was elevated. (The green p-value represents the difference of both repair types compared to native cartilage, and grey p-value represents the difference between treatment groups)

We further performed Raman spectroscopy to examine the biochemical composition of tissue formed in the repair regions compared to the surrounding articular cartilage, especially highlighting the major collagen and proteoglycan molecules (**Figure 5C**). Collagen content in gECM hydrogels was measured as 0.254 (normalized peak intensity), greater than the collagen content in microdrilling tissue (0.240; p = 0.0067). Both repair tissues showed greater collagen content compared to the surrounding articular cartilage (p=0.016). Relative proteoglycan content was increased in the microdrilling repair tissue compared to the surrounding articular cartilage and gECM hydrogels (p=0.017), while the proteoglycan content of gECM hydrogel was not different from native tissue. We additionally determined the proteoglycan-to-collagen ratio, a critical metric that indicates relative contributions to the macromolecular network governing the compression and swelling of cartilage tissue during movement (*25, 26*). The ratio in microdrilling repair tissue was greater compared to both native tissue (p=0.0136) and gECM hydrogels (p=0.043), indicating that microfracture did not maintain the critical structural and compositional balance of proteoglycans and collagens.

### Repair tissue maintains gross structure, evidenced by T2, T2*, and T1rho measurements similar to native cartilage

MRI analysis was used to further investigate the morphology of the repair tissue and complement the compositional measurements from immunohistochemistry and Raman spectroscopy. Both repair types showed similar tissue morphology, visualized using noninvasive MRI (**Figure 6A**). We measured variable quantitative MRI (relaxometry) measures, but mean values were not different compared to native cartilage based on T2, T2*, and T1rho (p>0.05) (**Figure 6B**). For T2, which correlates to water content and collagen alignment, gECM hydrogel repair was more variable and elevated compared to native cartilage, suggesting increased hydration. For T2*, which correlates to collagen structure and orientation, the microdrilling repair displayed lower values compared to native cartilage, suggesting lower water content, higher collagen density, and tighter collagen organization, typical of a fibrous repair. For T1rho, which correlates to proteoglycan content, neither repair type differed from native cartilage.

**Figure 6.**
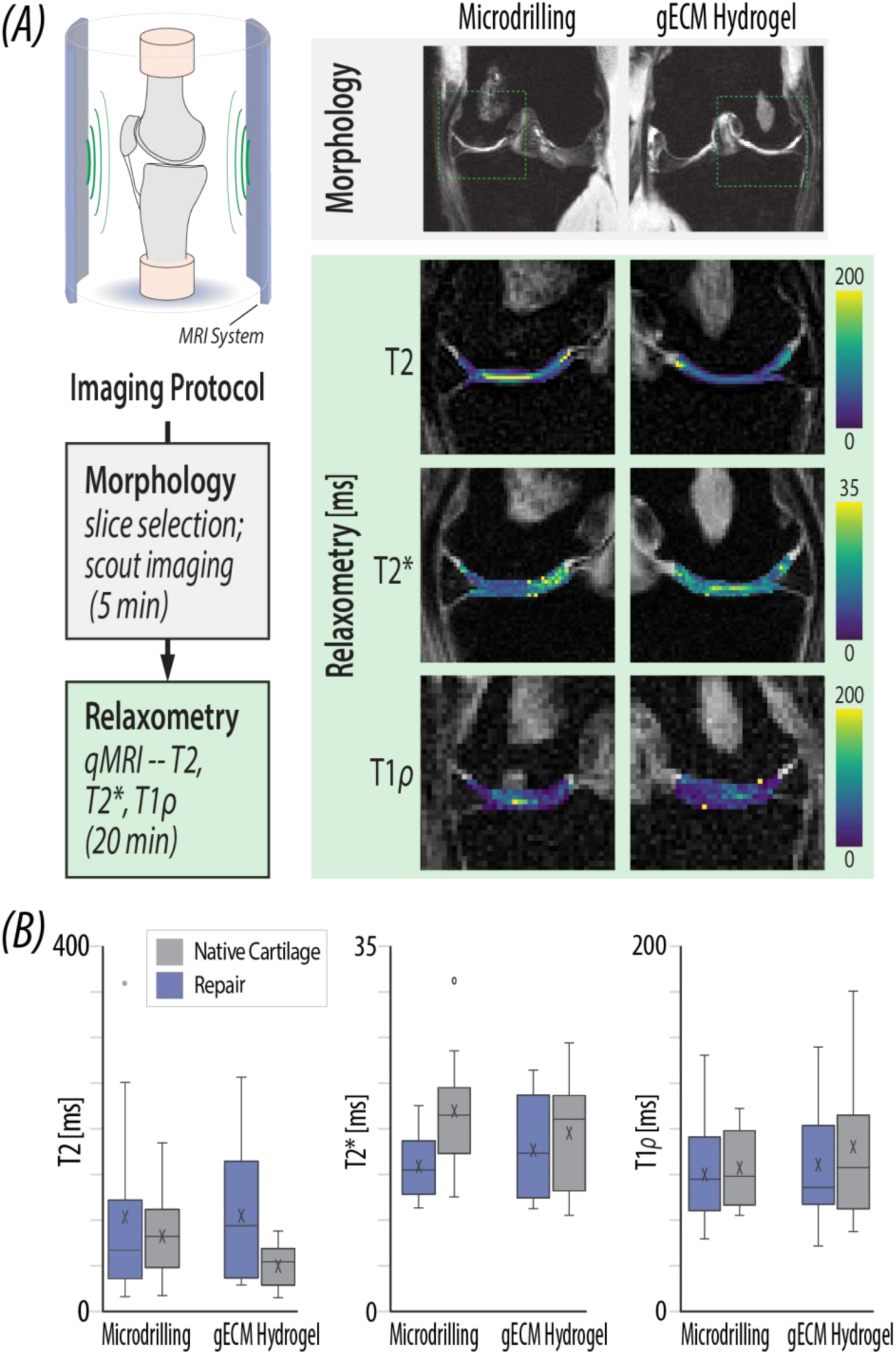
Repair tissue maintains gross morphology and structure, evidenced by T2, T2*, and T1rho measurements similar to native cartilage. *(A)* Schematic setup of the experiments. Morphological images show a cross section of the condyle where the defects were made and repaired with either gECM hydrogel or microdrilling only. Relaxometry metrics were evaluated in the load bearing region of the condyle including both the repair and surrounding native tissue. *(B)* Relaxometry analysis indicates biochemical similarities between the native (gray) and repair tissue (blue), in all repair groups, as evidenced by T2, T2*, and T1rho mapping.

### Repair tissue contains lower bone volume and trabecular thickness than native tissue

Finally, we evaluated the subchondral bone quality beneath the repair tissues using micro computed tomography (**Figure 7**). Bone density, a measurement of bone volume/tissue volume, was similar in both repair groups and was significantly lower than native tissue (p<0.0001). Trabecular spacing, or the average distance between individual trabecula, was similar between both repairs and native tissue. Finally, trabecular thickness was significantly lower in the mineralized portions of both repair types compared to native bone (p<0.0001). Because bone volume and quality typically depend on mechanical loading during normal activities (e.g., gait) and repair tissue surfaces were slightly recessed (gECM hydrogels) or absent (microdrilling) during delivery, decreased density and trabecular thickness are consistent with lower force transfer to repair tissues *in vivo*.

**Figure 7.**
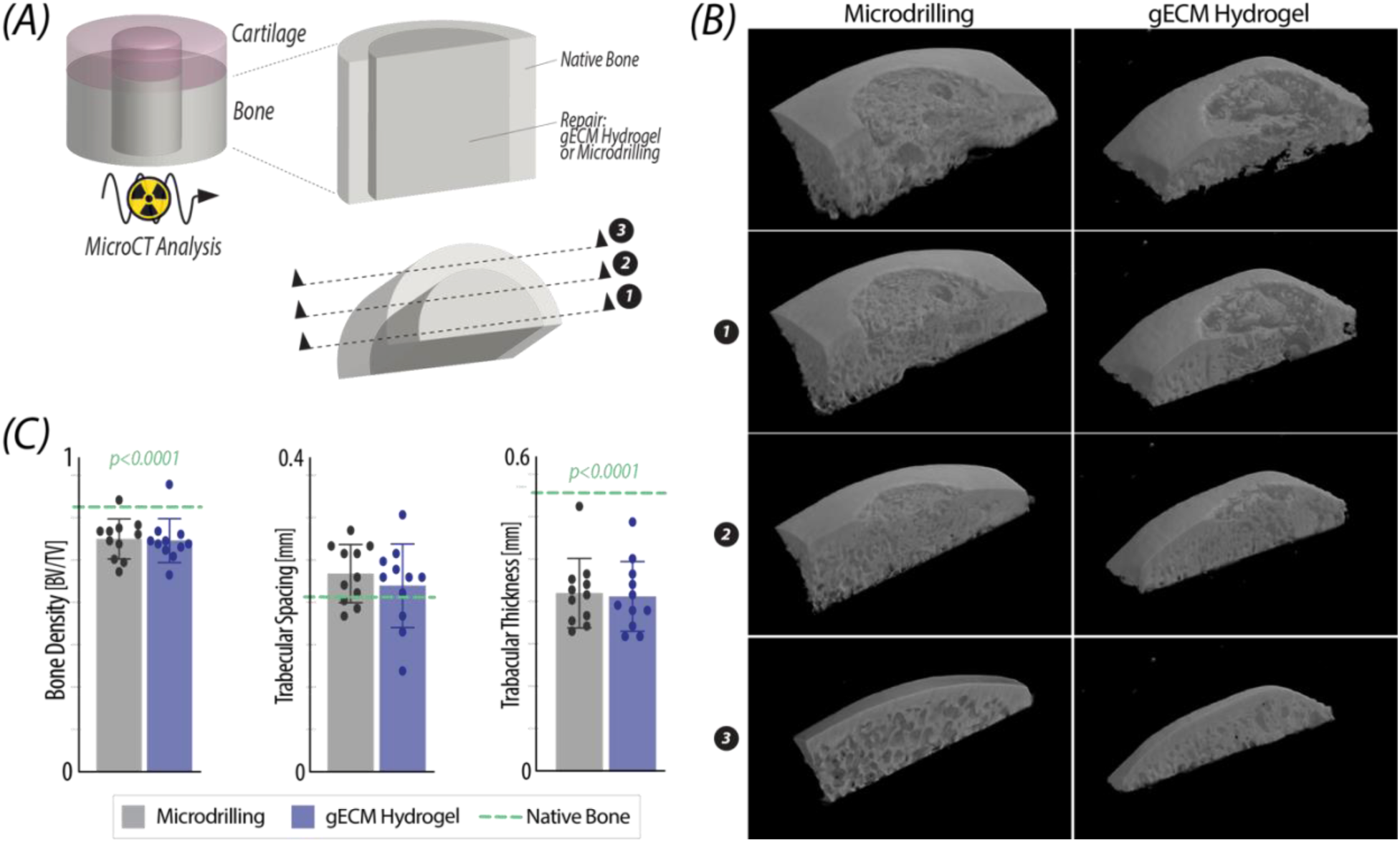
Repair tissue contains lower bone volume and trabecular thickness than native tissue. *(A)* Radiographic (micro computed tomography; microCT) analysis was used to visualize and quantify bone regions of the native tissue and repair tissue. *(B)* Reconstructed microCT scans of each type of repair shown from the center of the bisected plane (1) towards the outside edges of the implant (3). *(C)* Quantification of the bone volume/tissue volume (bone density), the average trabecular spacing, and the average trabecular thickness indicate that repair tissues contains lower bone tissue volume and trabecular thickness compared to native tissue (represented with dashed green line, ****p < 0.0001).

## DISCUSSION

Benchtop studies have demonstrated the potential of gECM hydrogels to mimic the structure and function of a tissue without limiting cellular invasion into the hydrogel following repair (*6, 17, 27*). Here, we have advanced and translated a gECM hydrogel for on-demand and flowable delivery to a critical-sized cartilage defect in the load-bearing region of a goat knee. The granules in the hydrogel were processed to remove cellular content but maintained many of the critical structural proteins in cartilage, including collagens, proteoglycans, and a myriad of ECM-related proteins. The ability to form gECM hydrogels using thiol-modified hyaluronan motivated our desire to explore translational studies in a large animal model. Accordingly, after 12 months *in vivo*, >80% of the defects in goat knee cartilage were filled following both microdrilling and ECM hydrogels. Further analysis enabled characterization of the content and function of the tissue in each repair type. We have shown that the gECM hydrogel, compared against microdrilling as a clinical standard of care, additionally displayed low surface friction, adhesion, and roughness to enable smooth sliding within the joint. The tissue within the gECM hydrogel repair restored the proteoglycan:collagen ratio measured in native tissue, and the type of collagen in the gECM hydrogel repairs was predominantly type II, rather than the majority type I observed in microdrilling repair. Functional MRI showed that the gECM hydrogel repair maintained some of the native tissue structure, with similar T2* and T1rho measurements between native tissue and repair. However, we also found that several metrics do not differ significantly between microdrilling and gECM hydrogel repairs, but did differ significantly in both groups from native tissue. Contact modulus, equilibrium modulus, bone tissue volume and trabecular size, and T2 MRI quantification (representative of tissue hydration), all demonstrate a quality of repair tissue that does not fully restore the native cartilage properties by 12 months *in vivo*. It is further possible that these parameters may approach native cartilage values with additional healing time.

gECM materials as flowable, tissue-engineered biomaterials show promise in the treatment of articular cartilage defects. An ideal material for cartilage repair must restore the structural architecture that is essential to cartilage function, as the tissue experiences routine loading of multiple times body weight (*28*). Ideally, the repair material would be able to withstand this loading shortly after delivery to the defect site to enable a quick recovery back to load bearing. However, it is also critical that the repair enables cellular migration and new matrix deposition into the defect space to ensure long term integration with the surrounding tissue environment. Previous *in vitro* studies and subcutaneous mouse models showed that pulverizing ECM and reconstituting the particles in a chondrogenic hydrogel supported cellular migration while maintaining mechanical properties of a typical osteochondral tissue (*6*). However, one of the key challenges facing current clinical repair solutions for cartilage is that scaffolds do not integrate into surrounding tissue and ultimately fail to provide a long-term (>1 year) repair (*29*). Consequently, long-term (>6 month) evaluation of new materials in physiologically-relevant loading environments, such as large-animal models, is essential to improve translational success.

Goats are a well-established large-animal model for cartilage repair due to their attainment of skeletal maturity by 2–3 years of age, sufficiently thick articular cartilage for defect creation (0.8–2 mm), and lack of spontaneous healing in critical-sized defects (>6 mm diameter) (*30, 31*). However, disruption of the stiff subchondral bone plate in this model has been associated with cyst formation (*32*). In our MRI analyses, we observed localized signal changes in a subset of treated joints, consistent with previously reported responses to microdrilling in goats (*30*). Species-specific differences must also be considered when translating results, particularly the fact that humans are bipedal whereas goats are quadrupedal, resulting in higher joint loads in humans. The granular design of the gECM hydrogel, which has previously been shown to exhibit increased compressive modulus under higher loading and was confirmed to be mechanically comparable in pre-loaded testing in this study (**Figure 1**), may be well suited to address this translational challenge (*33*). Building on prior *in vitro* evaluations, this study assessed the performance of gECM hydrogels in a goat model over 12 months as a critical step toward clinical translation to humans.

Successful clinical translation of gECM hydrogels depends not only on long-term functional outcomes, but also on whether new therapies meet the practical and economic constraints of orthopedic surgery. Currently, the most widely used cartilage repair options are osteochondral allografts and matrix-induced autologous chondrocyte implantation (MACI), yet both have substantial limitations. Osteochondral allografts can provide durable pain relief and functional improvement (*34, 35*), but their use is restricted by limited donor availability, long wait times, and high cost, which is often not fully reimbursed. MACI, a two-stage, cell-based therapy, is similarly constrained by high cost (>$35,000), limited patient eligibility due to joint degeneration, potential de-diferentiation of chondrocytes prior to implantation, and the need for open surgical procedures. As a result, broad adoption of either approach remains limited.

Minced cartilage–based therapies have emerged as a promising alternative (*36–40*) with commercial products such as DeNovo® NT and BioCartilage®. However, these solutions depend on scarce and short shelf-life juvenile cartilage and/or non-physiologic fibrin glue to secure minced tissue matrix within the defect, and can require complex, multi-step surgical techniques that reduce ease of use and reproducibility. Moreover, limited preclinical and clinical evidence has further slowed their adoption. These challenges highlight key design considerations for translational cartilage repair materials. The gECM hydrogel evaluated here addresses several of these limitations by using nonviable minced tissue crosslinked with a natural hyaluronan matrix, eliminating the need for fibrin glue while providing immediate implant fixation in a single-stage procedure. The gECM hydrogel design improves surgical efficiency, supports earlier load bearing, and enables an off-the-shelf, cost-effective therapy with the potential for broader clinical adoption.

This study employed porcine-derived cartilage particles to generate the acellular granules used in the gECM hydrogels; however, clinical translation requires careful consideration of tissue sources and regulatory pathways. Although decellularization reduces immunogenic risk, porcine-derived materials may introduce allergy concerns and increase regulatory burden. Transitioning to a human cartilage source could mitigate these risks while enabling a more clinically relevant, off-the-shelf, allograft-like material. Notably, substantial volumes of high-quality human cartilage are currently discarded due to the geometric constraints of osteochondral allografts, and the gECM platform could repurpose this tissue (i.e., extend the “gift of donation”) without size or shape limitations. However, incorporation of cartilage particles into a thiol-modified hyaluronan matrix exceeds minimal manipulation, shifting regulatory classification away from transplant tissue alone. Accordingly, successful translation will depend on future work that establishes scalable manufacturing, batch-to-batch consistency, and safety in alignment with regulatory expectations.

An unexpected finding of this study was the strong performance of the microdrilling group, which exceeded the expected preclinical efficacy. In clinical practice, many surgeons have moved away from techniques like microfracture due to inconsistent outcomes and poor long-term repair (*41–43*). However, the procedure used here differed from standard clinical microfracture; due to veterinary constraints and the hardness of goat subchondral bone (*31*), we instead used a microdrilling technique involving multiple 1-mm diameter holes drilled to approximately 5-mm depth. This approach may have enhanced marrow access, progenitor cell recruitment, and defect filling. Notably, prior work has shown that microdrilling techniques can outperform traditional microfracture approaches in tissue repair (*44*). Consistent with this, we observed >80% defect fill across both repair groups, whereas clinical microfracture in humans is often associated with incomplete or uneven filling dominated by fibrocartilage (*41*). Given these factors, and current recommendations for large-animal studies (*31*), repair efficacy was evaluated relative to native cartilage rather than as a fold change from the microdrilling repair tissue.

The gECM hydrogels tested here show promise in restoring several native cartilage properties, including critical surface (tribological) parameters, but several limitations should be acknowledged. First, a defining measure of clinical success is alleviation of pain and improvement in quality of life, which are difficult to assess in goats except through severe gait changes; formal gait analysis at baseline and post-surgery would have strengthened the study. Radiographs were used longitudinally over 12 months but, while effective for detecting gross joint changes, they are limited in resolving subtle cartilage (soft tissue) repair. Imaging with MRI or arthroscopy at intermediate timepoints (e.g., 6 or 9 months) could have provided more detailed evaluation. After sacrifice, repaired and surrounding tissue were harvested for structural and compositional analyses, but the rough cartilage surface introduced variability in indentation measurements. This was addressed by fitting the Hertzian contact modulus over a controlled range of indentation depths, though surface irregularities still influenced measurements of fluid load fraction.

In this study, we evaluated a potential clinical therapy for critical-sized defects in the load-bearing region of the knee condyle using a goat model. Defects were filled with gECM hydrogel during surgery, and animals returned to normal activity for one year. The study represents a key translational step, as material preparation and application closely mirrored anticipated human surgical workflows. These results support a shift from cell-based or multi-stage procedures toward a single-stage, off-the-shelf biomaterial capable of immediate and durable load-bearing repair. The demonstrated *in vivo* performance provides strong evidence that gECM hydrogels can meet the mechanical, biological, and surgical requirements for articular cartilage repair, positioning this approach as a promising paradigm for clinical translation.

## MATERIALS AND METHODS

### Study design

The objective of this study was to evaluate the therapeutic potential of a unique gECM biomaterial in a critical-sized defect in the knee cartilage joint of a goat. The study was intentionally designed to evaluate the material’s repair efficacy over a 12-month period to evaluate long-term repair potential. We hypothesized that the gECM biomaterial would integrate with surrounding tissue functionally and mechanically, enable cell migration throughout the repair, and replicate the critical cartilage surface parameters. The *in vivo* goat study was approved by the Colorado State University Institutional Animal Care and Use Committee (IACUC) and performed as directed by the IACUC. gECM biomaterials were implanted in the femoral condyle of female goats with no other fixation for 52 weeks (n=11).

### Granular ECM hydrogel sourcing, processing, and manufacturing

The gECM hydrogel consists of 10mg/mL hyaluronan (18-21% thiolation) packed with 0.2g/mL cartilage microparticles, mixed at the point of care in the surgical suite.

#### Component A

Cartilage tissue was sourced from porcine animals within 48 h of slaughter. Tissues were pulverized using a liquid nitrogen cryo-mill and sorted via a micro-sieve stack to isolate microparticles 40-100 μm in diameter (Electron Microscopy Sciences, Hatfield PA). Cartilage microparticles were decellularized in 2% sodium dodecyl sulfate (SDS) for 8 h at 37°C, and in 0.1% DNase for 3h to remove xenogeneic cellular and genetic material. Microparticles were flash frozen in liquid nitrogen, and lyophilized. The particles were mixed with 500 µl of sterile PBS to create a slurry of 0.4g/mL and loaded in a sterile syringe.

#### Component B

Glucoronate carboxyl groups on hyaluronan were replaced with thiol groups following previously established protocols (*45*). Briefly, hyaluronan (MW 100 kDa, Lifecore Biomedical) was dissolved at 10 mg/ml in degassed milliQ water. Dithiobis propanoic dihydrazide (DTP) (Frontier Scientific) was added to the solution, the pH was lowered between 4.5 and 4.75, and then (1-ethyl-3-(3-dimethylaminopropyl) carbodiimide (EDC) (ThermoFisher) was added to begin the reaction. The pH was maintained between 4.5 and 5 for 50 minutes. After 50 minutes, the reaction was stopped by raising the pH above 7. Next, dithiotreitol (DTT) (Fisher Scientific) was added, the pH was raised above 8.5, and the solution was stirred for 24 h at room temperature. After 24 h, the pH of the solution was lowered, the liquid was transferred to dialysis tubing (10 kDa membrane cutoff, Spectrum Labs) and the solution was dialyzed in an HCl solution supplemented with 100 mM sodium chloride for 8 solution changes, then in an HCl solution without supplements for 4 solution changes. Dialysis was completed in a sealed chamber with continuous nitrogen gas bubbling into the HCl solution. With each batch of HA thiolation, the substitution rate was confirmed to be 18-21% (thiolated mmols/unthiolated mmols) using a standard Ellman’s assay following manufacturer’s protocol (Ellman’s solution, ThermoFisher).

### DNA quantification in tissue matrix

Using a Sigma Aldrich Mammalian Genomic DNA Miniprep kit (GenElute™ Mammalian Genomic DNA Kit) and specified protocol, DNA was extracted and the concentration was analyzed. Briefly, the tissue was digested in the provided lysis solution with proteinase K at 55 C until no visible tissue remained (∼3 hours) and then spun through a DNA binding column. After several washes, the DNA was then eluted into a final solution for analysis. The ng/µL of DNA in the samples was calculated using a Thermo Scientific NanoDrop in absorbance mode at 260/280 nm, and normalized by dividing by the initial weight of the sample.

### Mass spectrometry of granules and native tissue

#### Sample preparation for proteomic analysis

Lyophilized, decellularized material from each sample (0.5-2 mg) was homogenized in 200 μL/mg freshly prepared hydroxylamin buffer (1 M NH_2_OH−HCl, 4.5 M Gnd−HCl, 0.2 M K_2_CO_3_, pH adjusted to 9.0 with NaOH). Samples were homogenized at power 8 for 3 minutes and incubated at 45°C with shaking (1000 rpm) for 4 h. Following incubation, the samples were spun for 15 min at 18,000 x g and the supernatant was removed and stored at −80°C until further proteolytic digestion. Extracted protein from each sample was subsequently subjected to enzymatic digestion overnight (16 h) at 37°C with trypsin (1:100 enzyme to protein ratio) using a filter aided sample preparation (FASP) approach as previously described (*46*) and desalted during Evotip loading.

#### LC-MS/MS analysis

Digested peptides (200 ng) were loaded onto individual Evotips following the manufacturer‘s protocol and separated on an Evosep One chromatography system (Evosep, Odense, Denmark) using a Pepsep column, (150 µm inter diameter, 15 cm) packed with ReproSil C18 1.9 µm, 120Å resin. Samples were analyzed using the instrument default “30 samples per day” LC gradient. The system was coupled to the timsTOF Pro mass spectrometer (Bruker Daltonics, Bremen, Germany) via the nano-electrospray ion source (Captive Spray, Bruker Daltonics). The mass spectrometer was operated in PASEF mode. The ramp time was set to 100 ms and 10 PASEF MS/MS scans per topN acquisition cycle were acquired. MS and MS/MS spectra were recorded from m/z 100 to 1700. The ion mobility was scanned from 0.7 to 1.50 Vs/cm^2^. Precursors for data-dependent acquisition were isolated within ± 1 Th and fragmented with an ion mobility-dependent collision energy, which was linearly increased from 20 to 59 eV in positive mode. Low-abundance precursor ions with an intensity above a threshold of 500 counts but below a target value of 20000 counts were repeatedly scheduled and otherwise dynamically excluded for 0.4 min.

#### Global proteomic data analysis

Data was searched using MSFragger v4.1 via FragPipe v21.1 (*47*). Precursor tolerance was set to ±15 ppm and fragment tolerance was set to ±0.08 Da. Data for porcine origin samples was searched against UniProt restricted to *Sus scrofa* with added common contaminant sequences (46,291 total sequences, downloaded 6/1/24). Enzyme cleavage was set to semi-specific trypsin for all samples. Fixed modifications were set as carbamidomethyl (C). Variable modifications were set as oxidation (M), oxidation (P) (hydroxyproline), deamidation (NQ), Gln->pyro-Glu (N-term Q), and acetyl (protein N-terminus). Label free quantification was performed using IonQuant v1.10.27 with match-between-runs enabled and default parameters. Results were filtered to 1% FDR at the peptide and protein level. Following database searching, proteins were annotated as cytosolic, nuclear, membrane, cytoskeletal (CS), or matrisome based on a concise list of cellular compartments derived from The Matrisome Project (*48*) and Gene Ontology (GO) enrichment terms (*49*). Matrisome proteins were categorized further as collagens, proteoglycans, ECM glycoproteins, secreted factors, ECM regulators, or ECM-affiliated proteins based on MatrisomeDB classifications (*50*). Any remaining uncategorized proteins were manually annotated with the aid of the Human Protein Atlas (*51*). Raw intensities were normalized such that the total summed intensity for each sample was equivalent and then log10-transformed.

### Bulk mechanics and rheological testing of gECM constructs

Components A and B were mixed and immediately tested with the rheometer (first 0.3 mL of combined gECM hydrogel), or molded into cylindrical PDMS constructs (remaining freshly prepared gECM hydrogel). Rheological shear sweeps (0.1 to 250 s^-1^) were performed at room temperature within 5 minutes of mixing using a rotational rheometer (Anton Paar). Cylindrical biomaterial constructs were molded and polymerized for 45 minutes at 37°C. Bulk mechanical testing was performed after polymerization and 12 h in PBS under unconfined compression to 40% strain at a quasi-static strain rate of 0.1%s^-1^ (Anton Paar), and the compressive modulus was calculated from the linear region between 30-40% strain.

### Animal selection and anesthesia

All study procedures were approved by the Colorado State University Institutional Animal Care and Use Committee and were performed in an AAALAC accredited facility. Eleven (n=11) healthy, skeletally mature conventionally raised Boer-cross does were enrolled in this study. All animals were fed a standard diet of alfalfa and grass hay mix with grain supplementation, as needed. Animals were co-housed indoors in box pens for a minimum of 2-weeks following surgical procedures, followed by a minimum of 2-weeks in an indoor/outdoor pen. Animals were then housed on dirt pasture with access to 3-sided shelters for the remainder of the study.

For surgical procedures, general anesthesia was induced by injecting a combination of midazolam (0.1 mg/kg) and ketamine (1.0-3.0 mg/kg) intravenously into a peripheral ear venous catheter. Anesthesia was maintained using isofluorane (1.5-3%) in 100% oxygen through an endotracheal tube. Blood pressure was monitored continuously throughout the procedures either through a peripheral arterial catheter or blood pressure cuff. One day prior to surgery, a transdermal fentanyl patch (50 mcg) was adhered to the skin on the axillary region of each animal for sustained release over five days and phenylbutazone (1 g) was administered once per day orally for seven days for analgesic effect. Additionally, a dose of tulathromycin (2 cc) was administered subcutaneously both the day prior to surgery and on the fifth day post-op for prevention of infection. Buprenorphine (0.005-0.01 mg/kg) was administered subcutaneously for post-op analgesia, as needed.

### Bilateral cartilage resurfacing of the medial condyle in goats

Each animal underwent a medial parapatellar mini-arthrotomy on both the left and right knees (bilaterally) to expose distal medial femoral condyles. A 6.5 mm drill guide was impacted into the central portion of each medial femoral condyle to a depth of approximately 2 mm. An end-mill with a stop was placed down the drill guide and used to create an osteochondral defect (6.5 mm Ø; 2 mm depth) through the cartilage and calcified cartilage layers in the central weight-bearing region of the medial femoral condyle for each (right and left) stifle. Saline irrigation was applied liberally to the end-mill during drilling. In the control microdrilling groups, a 1.1 mm drill bit was utilized to create five small microdrilling holes into the subchondral bone. The defect was left empty in the control groups. In the cartilage resurfacing groups, a similar microdrilling procedure was performed and the defect was then filled with the gECM hydrogel material to contour the surface of the cartilage. Following either microdrilling creation or condylar resurfacing (control vs. experimental), the synovial capsule was approximated with absorbable suture in an interrupted pattern. The fascia and any dissected muscle tissue was then closed using absorbable sutures in a continuous pattern, followed by closure of the subcutaneous layer. Finally, the skin was closed using a Ford-interlocking pattern with non-absorbable suture material.

### Radiographic imaging

All animals received radiographic imaging of the stifle joints at five study timepoints: immediately post-op, 13, 26, 39, and 52-weeks. Radiographic images were performed in the anteroposterior and lateral planes (voltage 80 kVp, current 1.0-1.5 mAs) at each time point using a Sound Fusion DR® radiography system (SN: FSN0041, Sound, Carlsbad, CA, USA) under minimal restraint. Radiographs at 52-weeks were performed as explanted, *en bloc* hindlimbs.

### Euthanasia and necropsy

At 52-weeks, all animals were humanely euthanized by intravenous overdose of pentobarbitone solution (88 mg/kg). A gross necropsy, including assessment of all major organs, was performed and both hindlimbs were harvested for further analysis.

### Whole-joint analysis of structure and mechanical function by relaxometric MRI

All joints with defects on the medial condyle were thawed at 4°C for 48 h before the experiment and wrapped with phosphate-buffered saline-soaked paper towels to prevent dehydration. The knee imaging protocol consisted of a fast gradient echo MR image sequence (scout) for localization and subsequently 3D double echo steady state (DESS) acquisition, quantitative T2, T2* and T1ρ measurements. All imaging was performed using a clinical MRI system (3 T; Siemens Prismafit) with a 15-channel knee coil (Tx/Rx Knee 15 Flare Coil, QED, LLC). The imaging settings used are listed in Table 1. All relaxometry methods employed fat suppression. For T2 and T2* maps, the slice registered to the same position with T1ρ was selected for analysis. The tibiofemoral contact area on the medial condyle was selected for the ROI and the spatial average (SA) of the pixels within the ROI was calculated.

**Table 1.**
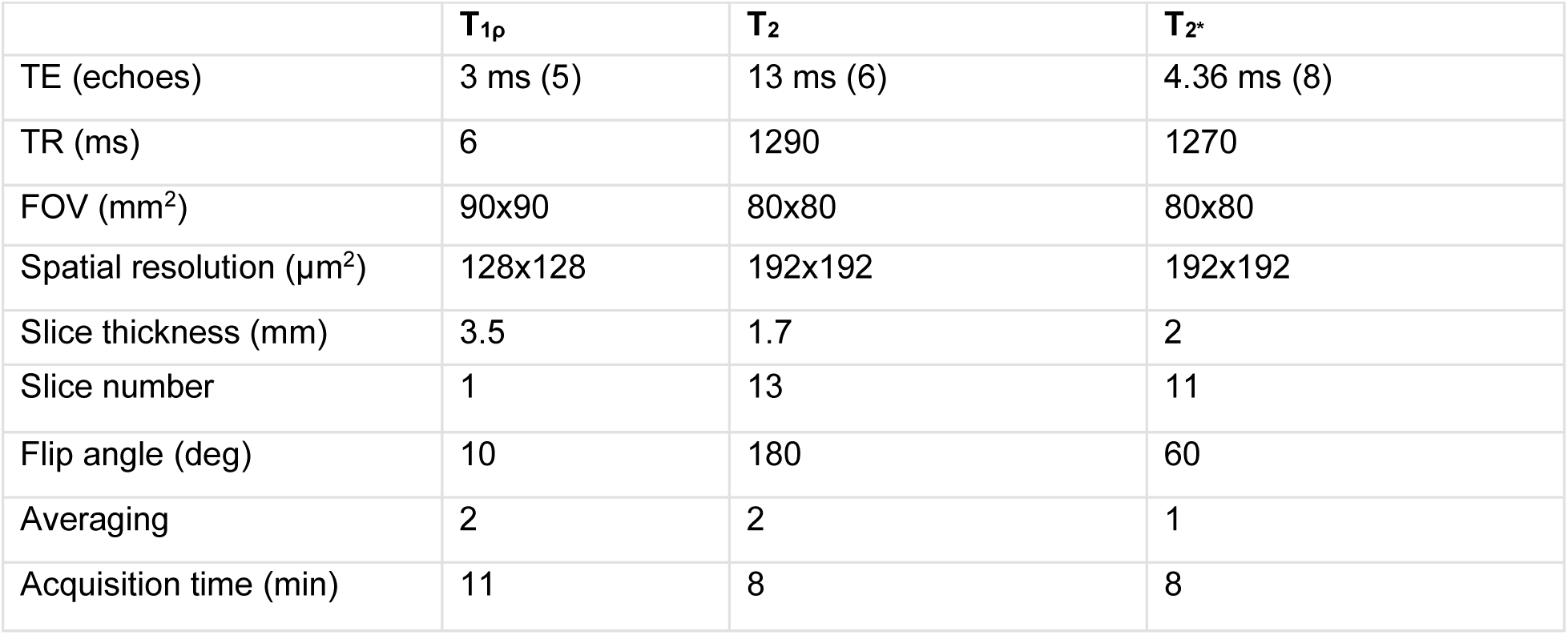
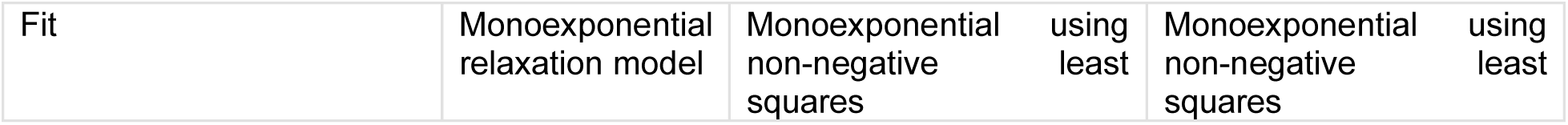
Qualitative MRI acquisition parameters.

### Repair tissue joint dissection and extraction

Following MRI analysis on the whole intact joints, the joints were opened, and 10 mm plugs extracted (including repair tissue with surrounding native tissue). Dissected tissue plugs with repair and native tissue were bisected with a sharp blade. One half of the tissue was fixed in formalin for bone XRM and histological analysis and the other half was flash frozen in liquid nitrogen for mechanical analysis.

### Quantitative imaging of osteochondral bone repair and remodeling

Trabecular bone volume fraction, trabecular thickness, and trabecular spacing were measured using micro-computed tomography (XRadia Versa 520, Zeiss, Dublin, CA, USA). The explants were scanned with a 0.4× objective, voxel size of 13.4 μm3, energy settings of 70 kV, 6.0 W, 1 s exposure, 801 projections, and using the LE2 filter. Automated centering and beam hardening corrections were applied using Scout-and-Scan Control System Reconstructor software (v 14.0.14829.38124). To analyze the bone structure, collected micro tomography images were imported into Dragonfly software (ORS v 4.1) and the Otsu algorithm (*52*) was implemented to separate ranges of the histogram corresponding to bone and background noise. Distinct volumes to analyze were designated in two key regions: native bone and tissue repair (i.e., microdrilling or gECM hydrogel). In each region, trabecular bone volume fraction was calculated. A sphere fitting-method was used in each trabeculae or in the space between trabecula to calculate trabecular bone volume fraction, trabecular bone thickness, and trabecular bone spacing using Dragonfly Bone Analysis (v2022). The filled-in bone volume in each region of interest was calculated using the Dragonfly Bone Analysis and compared to a total ideal bone volume to obtain bone tissue fill fraction.

### Structural histological staining and analysis

Plastic embedding, slicing, and staining were performed by Indiana University histology core. After formalin fixation, specimens are dehydrated through a graded series of ethanol (4 h per step), cleared in xylene (4 h) and infiltrated first with unpolymerized methyl methacrylate (4 h) and then for several days (3-7+) with unpolymerized methyl methacrylate containing 4% dibutyl phthalate (a softening agent). The specimens were then embedded in a medium of methyl methacrylate + 4% dibutyl phthalate + 0.25% Perkadox 16 (catalyst) and allowed to polymerize at room temperature. Excess plastic from the blocks was removed using a band saw and the blocks were shaped using a dental model trimmer. Thin sections (4 - 10 μm; primarily cancellous bone) were cut using a rotary microtome equipped with a tungsten-carbide knife.

Goldner’s Trichrome (GT): Sections were deplasticized using Xylene and re-hydrated (100%, 100%, 95%, 70%, H2O). Sections were stained in a working solution of Weigert’s Iron Hematoxylin for 7.5 minutes (*53*) followed by two sequential rinsing steps. Next, slides were stained in Phospomolbydic Acid – Orange G for 8 minutes, rinsed, and then stained in Light Green SF Yellowish for 15 minutes. Finally, slides were rinsed, dehydrated, cleared in xylene, and coverslipped with Eukit.

Toluidine Blue (TolB): Sections were deplasticized in acetone and re-hydrated (100%, 100%, 95%, 70%, H2O). Sections were etched in 0.1% Formic Acid for 60 seconds without agitation, rinsed in distilled water, and stained in Toluidine Blue for 3 minutes without agitation. Following staining, sections were rinsed, dehydrated and cover-slipped with xylene-based mounting media. Analysis: Image J analysis of GT -stained slides was used to calculate the percentage of the defect area that had been filled with some kind of tissue repair. Image J was used to create a mask of the tissue within the defect and was compared to the total volume of the tissue if the cartilage surface contour were completed by matching native tissue to native tissue on each side of the defect. Additional analysis was completed on the histology through standard scoring criteria: three independent and blind researchers evaluated both GT and TolB-stained slides to score 9 criteria of cartilage repair outlined by the International Cartilage Repair Society (ICRS) II scoring system.

### Immunohistochemistry for types I and II collagen

The locations of types I and II collagen were visualized using histological sectioning and immunostaining performed by Premier Laboratory (Longmont, CO). Detection of type I collagen by immunohistochemistry (IHC) was carried out using a rabbit monoclonal antibody directed against type I collagen from rabbit (EPR7785 from Abcam). Detection of α-collagen II by immunohistochemistry (IHC) was carried out using a rabbit polyclonal antibody directed against α-collagen II from rabbit (203001 from MD Bioproducts). Formalin-fixed paraffin-embedded (FFPE) samples were sectioned at 5µm and mounted onto charged slides. Slides were dried overnight, baked at 60°C, deparaffinized in xylene, rinsed in alcohol, rehydrated in water, and equilibrated in wash buffer (TRIS-buffered saline with 0.05% (v/v) Tween 20; Dako, K8007). Slides were then loaded onto an AutostainerLink48 stainer (Dako) and the remaining steps were carried out at room temperature. Within the autostainer slides were incubated in 3.0% (v/v) hydrogen peroxide for 5 minutes, followed by proteolytic induced epitope retrieval (PIER) via Proteinase K for 5 minutes (Dako, S3020) for type I collagen staining. For type II collagen, within the autostainer slides were incubated in 3.0% (v/v) hydrogen peroxide for 5 minutes, followed by Chondroitinase ABC (from Proteus vulgaris from Sigma C3667-10UN). Next a serum-free protein block was applied for 5 minutes (Dako, X0909), anti-Collagen I primary antibody for 30 minutes (1.09µg/ml) or anti- αCollagen II primary antibody for 30 minutes (1.09µg/ml), EnVision+ anti-Rabbit Labelled Polymer-HRP for 30 minutes (Dako, K4011), and DAB+ Chromogen Solution for 5 minutes (Dako, K4011). Wash buffer rinses were applied between appropriate reagents. Slides were then manually rinsed in tap water and counterstained for 6 minutes in an Automation hematoxylin (Dako, S3301). The slides were again rinsed in tap water, and wash buffer was used as the hematoxylin bluing reagent. The slides were then dehydrated in absolute alcohol solutions, cleared in xylene, and cover slipped.

### Compositional analysis of cartilage repair using Raman spectroscopy

Raman spectroscopy was used to quantitatively evaluate the biochemical composition of repair regions and the surrounding articular cartilage surface. Samples were first imaged using a 5× objective to delineate repair regions and surrounding tissue. Samples were submerged in PBS, and Raman spectra (Renishaw InVia, 785 nm laser) were collected at sites in a 4×4 array (175 µm spacing) using a 63× immersion objective for Raman shifts ranging from 700 – 1700 cm^-1^. At each point in the array, 8 accumulations of 6s exposure were taken. For each spectrum, cosmic rays were removed, a linear baseline was subtracted, intensity was normalized, signal-to-noise (SNR) ratio was calculated (excluding spectra with SNR < 15), and spectra were smoothed (*54*). The remaining spectral acquisitions were averaged and the following analyses were performed: relative proteoglycan content was calculated as the ratio between 1377 cm^-1^ / 1245 cm^-1^, collagen content was reported as the sum of the following normalized peak intensities: 814 + 855 + 919 + 1245 + 1271 cm^-1^, and proteoglycan to collagen ratio was measured as the ratio between chondroitin sulfate peaks (1062 + 1340 + 1377 cm^-1^) and collagen content (*54, 55*).

### Characterization of bulk mechanics using indentation

Microindentation was used to evaluate the tissue-level mechanical response within both the repair regions and neighboring uninjured articular cartilage. Articular cartilage surfaces were first visualized under a 50× objective to select indentation sites. Samples were submerged in phosphate buffered saline (PBS) during testing. Microindentation (Hysitron TI 950 Triboindenter, xZ-500 extended displacement stage; 250 µm conicospherical probe) was performed spanning 4×4 indent arrays, with 175 µm spacing in x- and y- directions. Each indent first applied a 20 µN preload for surface detection and initial surface contact was identified. A 40 µm lift-off ensured disengagement from the sample surface, and indentation was performed to a prescribed depth of 12.5 µm beyond the initial surface detection (trapezoidal displacement-hold: 15 µm/s loading and 45s hold at max displacement to permit force relaxation) (*55*). Following each test, the contact point was identified using the virtual contact point method described in (*56*). The reduced Hertzian contact modulus (*E_c_*; assuming ⫁=0.5) was obtained by fitting the Hertz sphere-half-space model to the loading response over post-contact depths > 7.5 μm (*57–59*), and then evaluating:

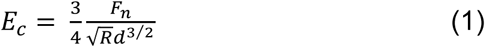

The fluid load fraction borne by tissue during indentation was calculated as:

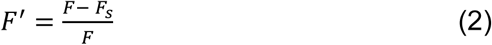

where *F_n_*is the normal force during loading, *R* is the sphere radius, *d* is the post-contact indentation depth, *F* is the total measured force, and *F_s_*is the force contribution from the matrix, estimated as described previously (*57*).

### Surface characterization by atomic force microscopy

Surface roughness, frictional coefficient, adhesion, compressive modulus, and topographical structures were measured using atomic force microscopy (Keysight 5500 AFM, Keysight Technologies Inc., Santa Rosa, CA, USA). All measurements were taken on the articular surface of the implant in the two regions of interest: native cartilage or tissue repair. A cantilever with known geometry was used (2 µm borosilicate sphere, NovaScan), and the cantilever stiffness was pre-calibrated to 0.36 N/m by the thermal fluctuation method (*60*). Lateral calibration was determined using an improved wedge calibration method (*61*) and a TGF11 silicon calibration grating (Mikromasch). Each sample was affixed to a μ-Dish high grid-50 dish (Ibidi, USA) using a viscous cyanoacrylate and hydrated for all AFM testing. At each location, the macroscopic region of interest on the articular cartilage surface was approached and a brief scan (4×4) was performed to ensure the height change was in an acceptable range (∼7 μm). After locating a measurable 40×40 μm area, sixteen independent locations of the scan area (4×4) were measured in force-volume mode, indented at 5 μm/s with a setpoint force of approximately 12 nN. To estimate the compressive modulus, force-displacement approach curves were fit to the Hertzian linear elastic model for a defined round tip geometry (*62, 63*). Adhesion force was measured as the pull-off force upon tip separation from the surface during probe retraction. Both adhesion and compressive modulus were extracted from force-distance curves using PicoView 1.14 AFM analysis software. Immediately following and within the same scan area, a high resolution contact scan (1028×1028 px, or 40nm/px) was performed at 20 μm/s with a constant applied normal force of 20 nN to generate detailed topography, raw deflection, and lateral voltage trace/retrace signals of surface features. Areas where the controller overloaded the sample or did not interact with the surface were excluded. Topographical images were analyzed to calculate root-mean-square (RMS) surface roughness using Gwyddion 2.56 SPM analysis software (*64*). The average lateral voltage signal over each area was converted to friction force by multiplying by the lateral calibration constant and dividing by the normal force to yield the coefficient of friction.

### Cell quantification using histological images

Cellularity of all regions of interest was quantified in native and repair tissue using plastic-embedded tissue slices (preparation detailed above) stained with Goldner’s Trichrome (detailed above). Each region (native or tissue repair) was isolated in a high-resolution image and imaged in the middle of the repair region or at least 3 mm away from the repair border for the native region. Standard ImageJ thresholding was used to identify cells (blue/black spots) and quantify cell number/tissue area using Image J particle counting methods.

### Statistical analysis

To test our hypotheses that treatment (gECM hydrogel or microdrilling) influenced articular cartilage architecture and repair (contact modulus, equilibrium modulus, fluid load fraction, friction, surface roughness, adhesion, protein signatures, and bone composition) in vivo, mixed model analyses of variance (ANOVAs) were performed on generalized linear models with treatment, animal, tissue region (native or repair region) and implant location (left stifle joint or right stifle joint) as the co factors, and the resulting measurement variable as the response. All residuals were checked and non-normal distribution were identified. The data was logarithmic transformed for statistical analysis, which was required for the contact modulus and equilibrium modulus data to maintain normality in the residuals.

## Acknowledgments

The authors acknowledge the histology core at the University of Indiana Histology Core Facility for completing the sectioning and staining of the plastic-embedded sections. The authors thank Elizabeth Chlipala and Amber Moser from Premier Laboratory, LLC for the sectioning and immunohistochemical staining of types I and II collagen. Mass Spectrometry was performed by the Mass Spectrometry Proteomics Shared Resource Facility [RRID SCR_021988] at the University of Colorado Anschutz. Raman spectroscopy, micro-computed tomography, and indentation testing was performed at the Materials Instrumentation and Multimodal Imaging Core Facility [RRID SCR_019307] at the University of Colorado Boulder.

## Author contributions

Conceptualization: CPN, JEB, JE

Methodology: JEB, KPM, JH, LS, KB, EM, HZ, WL, MCM, DCG, SB, VF, BG, SES, JE, BJ, NE

Visualization: JEB, JH, TL

Funding acquisition: CPN, JE, BJ

Supervision: CPN, JE, VF

Writing – original draft: JEB, KPM, LS, TL

Writing – review & editing: All authors

## Data and materials availability

All data, code, and materials used in the analysis are available to any researcher for purposes of reproducing or extending the analysis. For use of the exact materials studied in the manuscript, a material transfer agreement (MTA) would be necessary. All data is available from the corresponding author upon request.

## SUPPLEMENTAL FIGURES

**Supplemental Figure 1.**
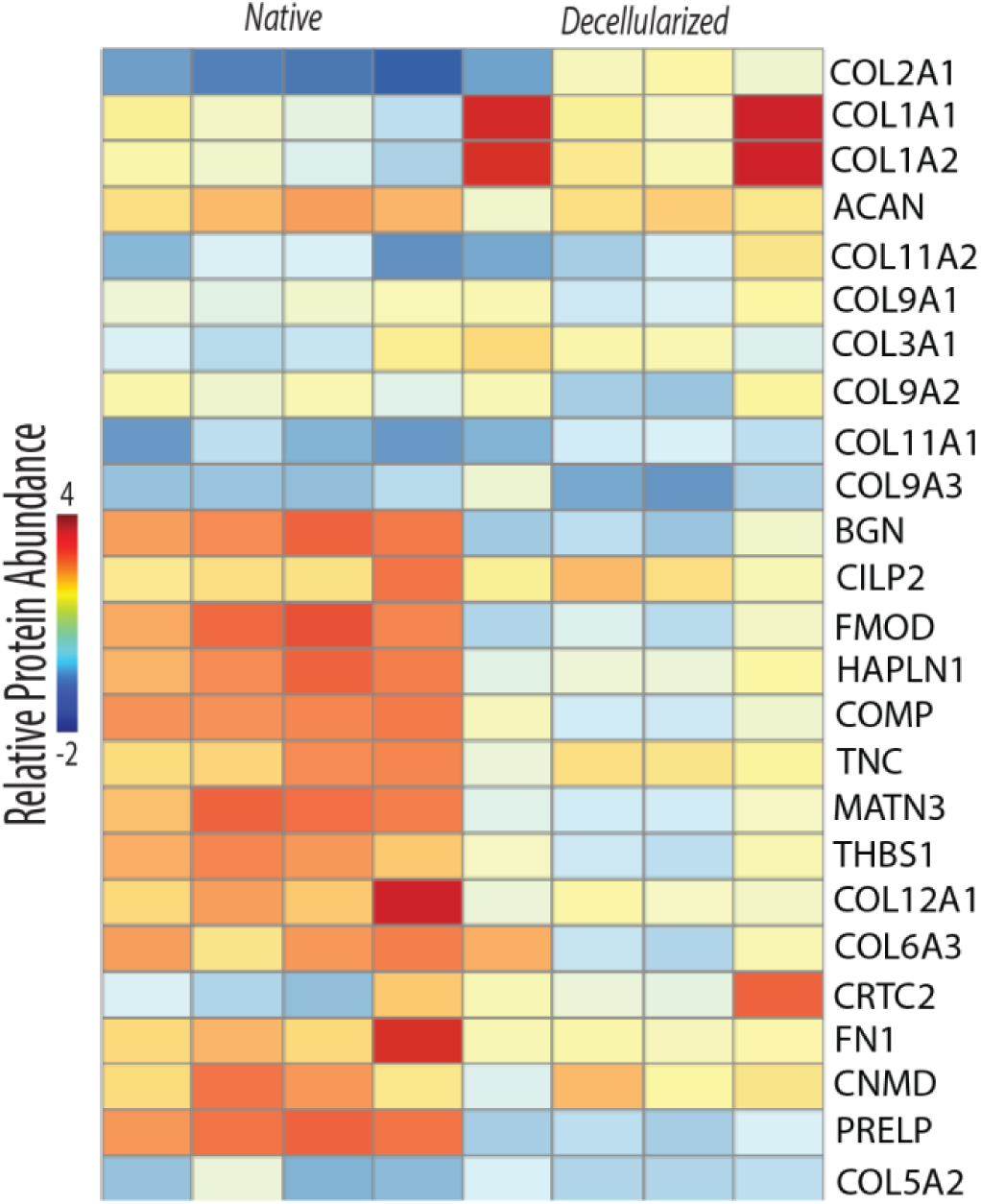
Proteomics analysis of the top 25 most abundant proteins by LFQ intensity. Collagens, aggrecan, and related common cartilage proteins were expressed in native and preserved in decellularized ECM granules.

**Supplemental Figure 2.**
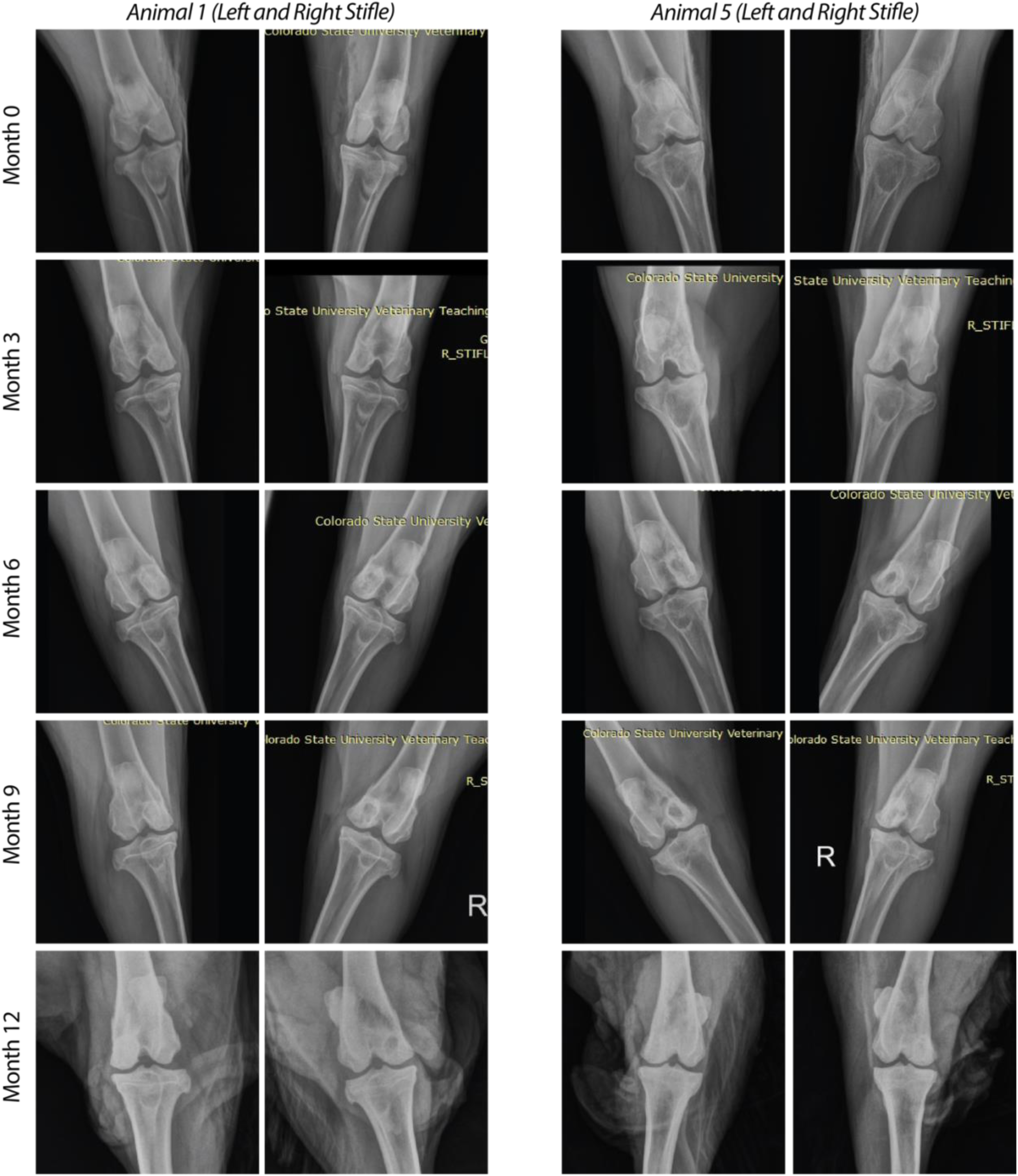
Timecourse radiographs of two representative animals, corresponding to histological data for animals in Figure 3. No evidence of joint space narrowing or other related bone abnormalities was detected.

**Supplemental Figure 3.**
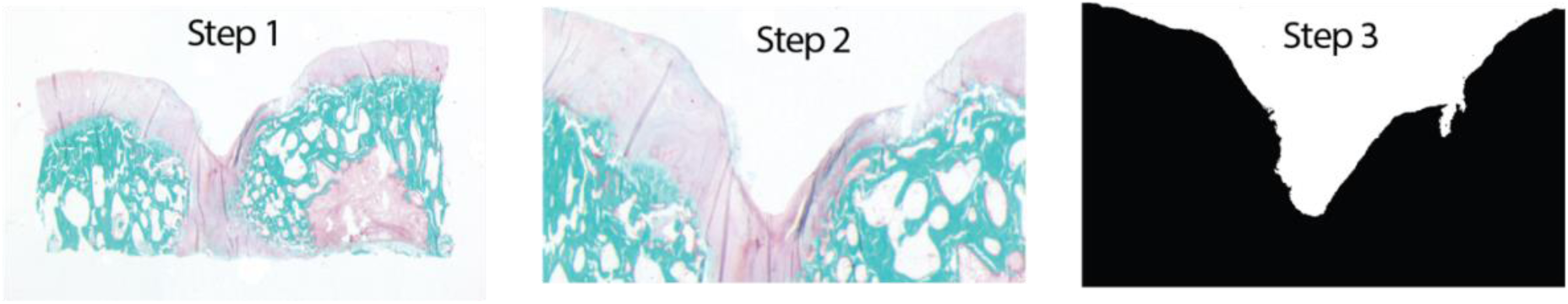
Method to process tissue filling. Images were acquired, cropped to consistent dimensions including the repair and surrounding cartilage regions, and thresholded. The percent area was computed from thresholded images as a measure of tissue fill.

**Supplemental Figure 4.**
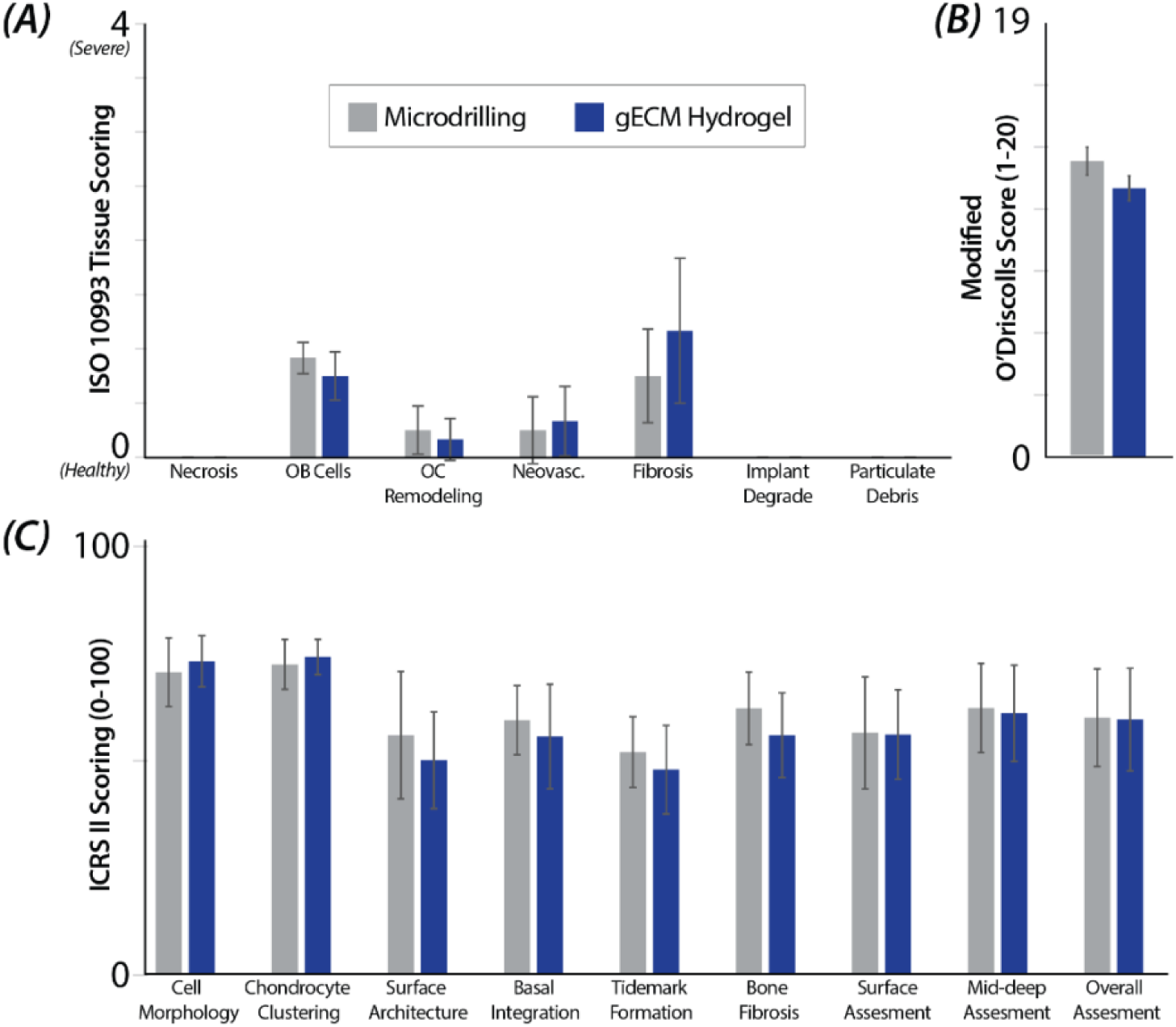
Scoring systems for cartilage repair. No differences in histology-based scoring metrics for microdrilling versus gECM hydrogels were detected. *(A)* ISO 10993 data shows metrics that were considered largely ‘healthy’. *(B-C)* ICRS II and Modified O’Driscolls scoring indicated similar responses for repair treatment, including specific metrics like surface architecture and overall assessment.

**Supplemental Figure 5.**
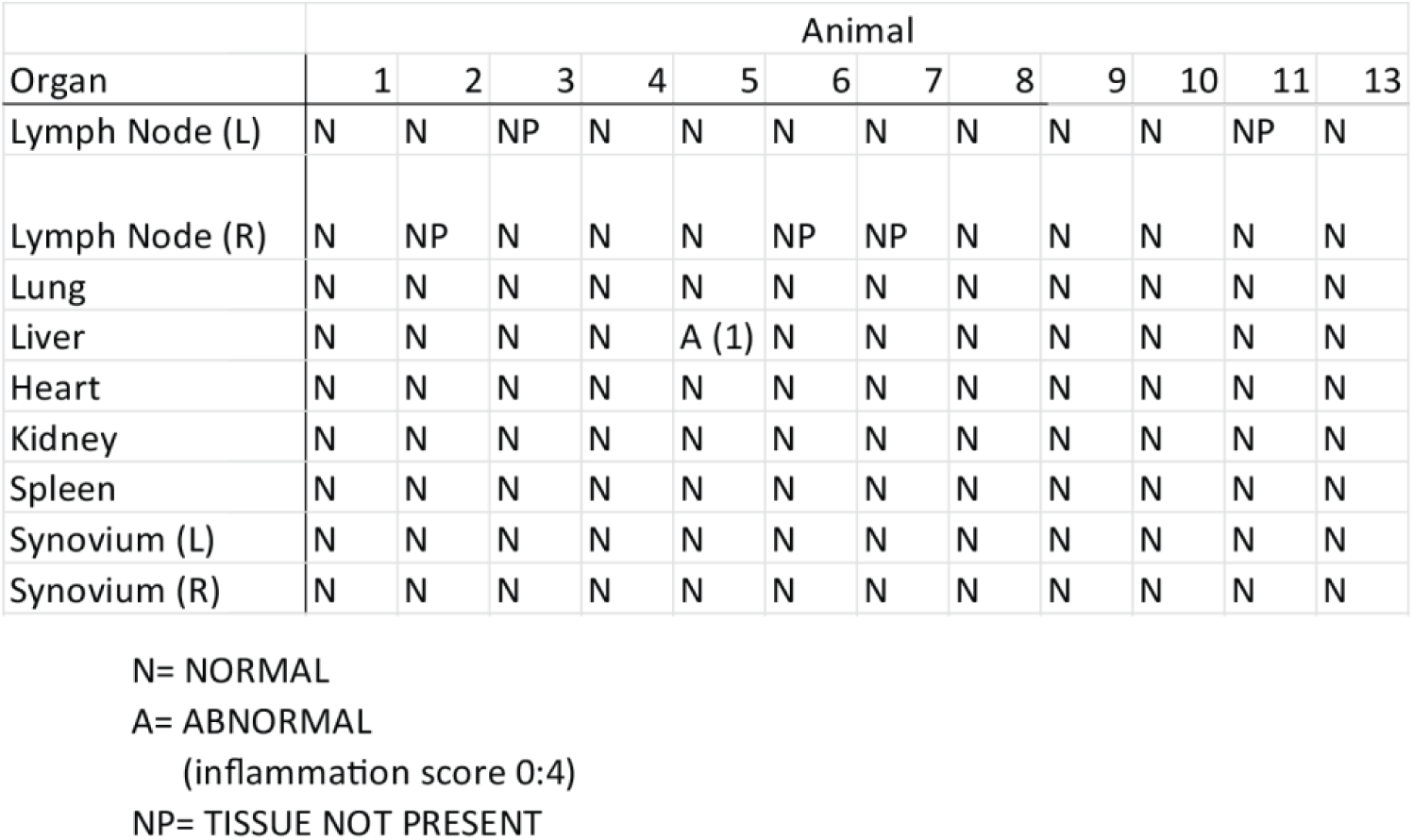
Gross systemic pathology for animals. Assessment of systemic pathology revealed largely normal appearance of lymph nodes, lung, liver, heart, kidney, spleen, and synovium.

